# Brain signal complexity and aperiodicity predict human corticospinal excitability

**DOI:** 10.1101/2024.02.09.579457

**Authors:** Joel Frohlich, Simon Ruch, Bettina H. Trunk, Marius Keute, Pedro A.M. Mediano, Alireza Gharabaghi

## Abstract

**Background:** Transcranial magnetic stimulation (TMS) holds promise for brain modulation with relevant scientific and therapeutic applications, but it is limited by response variability. Targeting state-dependent EEG features such as phase and power shows potential, but uncertainty remains about the suitable brain states.

**Objective:** This study evaluated broadband EEG measures (BEMs), including the aperiodic exponent (AE) and entropy measures (CTW, LZ), as alternatives to band-limited features, such as power and phase, for predicting corticospinal excitability (CSE).

**Methods:** TMS was delivered with randomly applied single pulses targeting the left primary motor cortex in 34 healthy participants while simultaneously recording EEG and EMG signals. Broadband and band-limited EEG features were evaluated for their ability to predict CSE using motor evoked potentials (MEPs) from the right extensor digitorum communis muscle as the outcome measure.

**Results:** BEMs (AE, CTW) significantly predicted CSE, comparable to beta-band power and phase, the most predictive and spatially specific band-limited markers of motor cortex CSE. Unlike these localized CSE markers at the site of stimulation, BEMs captured more global brain states and greater within-subject variability, indicating sensitivity to dynamic state changes. Notably, CTW was associated with high CSE, while AE was linked to low CSE.

**Conclusion:** This study reveals BEMs as robust predictors of CSE that circumvent challenges of band-limited EEG features, such as narrowband filtering and phase estimation. They may reflect more general markers of brain excitability. With their slower timescale and broader sensitivity, BEMs are promising biomarkers for state-dependent TMS applications, particularly in therapeutic contexts.

## Introduction

Therapeutic and scientific applications of transcranial magnetic stimulation (TMS) may be maximized by strategically timing stimuli to coincide with periods of increased corticospinal excitability (CSE), as indicated by cortical oscillations recorded through electroencephalography (EEG). However, uncertainty remains about the appropriate phase or frequency to target: Some studies report no significant EEG frequency or phase dependency of corticospinal excitability (CSE) (1), while others identify similar phases within either the alpha (2) or beta (3) frequency bands. Further studies have observed this relationship in both alpha and beta frequencies, though sometimes with opposing phases (4). Moreover, some findings suggest no direct phase or power dependency of CSE but reveal a phase-power interaction, where the phase dependency reverses direction with changing power levels (5). Importantly, methodological differences, including specific analytical approaches, may significantly influence the sensitivity to detect relationships between power, phase, and CSE (6) and may even lead to incorrect phase estimations (7).

To address the limitations of using EEG features based on phase or power within particular frequency bands as CSE markers, we explore different approaches. These include measures based on EEG signal entropy or “complexity”, and a measure based on the aperiodic exponent (AE) or “steepness” of the EEG 1/f background. In the former case, a broad literature of both theoretical (8–10) and empirical studies (11–14) suggest that cortical entropy, as measured using EEG or magnetoencephalography (MEG), relates to the cortical capacity for information processing, e.g., as indicated by states of wakeful consciousness (15) or lucid dreaming (16). Low entropy states often (17) − though not always (18, 19) − correspond to cortical hypersynchronization (e.g., slow waves), during which cortical circuits have fewer degrees of freedom and are thus less receptive to new inputs (12, 20). We therefore hypothesized that high motor cortical entropy, estimated using the Lempel-Ziv (LZ) and context tree weighting (CTW) algorithms, should predict high CSE. In the case of AE, evidence from a variety of experiments in different models (21), demonstrates that AE (i.e., 1/f steepness) increases with greater neural inhibition and decreases with greater neural excitation. As such, we hypothesized that low AE measured from motor cortical EEG signals should predict high CSE.

Both measures are computed from the broadband EEG signal and thus avoid the pitfalls of approaches which require the selection of a particular frequency band such as alpha or beta. Notably, these broadband EEG measures (BEMs) are also derived from longer EEG windows (1 s) than those used to estimate oscillatory phase or power, a fact which may reduce artifact susceptibility but also sensitivity to short fluctuations in CSE. In this sense, BEMs may be complimentary, rather than alternative, to conventional bandlimited EEG measures.

In what follows, we test our hypotheses using randomly applied single TMS pulses targeting the left primary motor cortex (M1) in young, healthy participants. During the experiment, EEG signals we recorded and later processed to compute bandlimited (alpha and beta phase and power) and broadband (LZ, CTW, and AE) EEG features. Electromyogram (EMG) signals from the right extensor digitorum communis (EDC) muscle were also recorded and used to compute motor evoked potentials (MEPs), with MEP amplitude serving as a readout of CSE. Our results suggest that both EEG entropy measures (LZ and CTW), as well as AE, are useful predictors of CSE and hold promise for guiding EEG-based TMS applications.

## Materials and Methods

### Data collection

We utilized available data from 36 healthy right-handed subjects (M = 24.5, SD = 3.25 years, 25 female), all of whom had no contraindications to TMS (22). All participants gave written informed consent, and the study was approved by the ethics committee of the Medical Faculty of the University of Tübingen.

TMS was applied using a MagPro-R30 stimulator with a MCF-B70 Butterfly Coil (MagVenture GmbH, Denmark) in combination with a neuronavigation system (Localite GmbH, Germany). Participants were seated in an armchair with their forearms pronated and fully relaxed. 64-channel EEG was recorded at 5000 Hz during stimulation (Braincap for TMS, Easycap GmbH, Germany; Brain Products DC amplifiers, BrainProducts GmbH, Germany). EMG of the right EDC was recorded at 5000 Hz with a BrainProducts ExG amplifier (BrainProducts GmbH, Germany) and Ag/AgCI AmbuNeuroline 720 wet gel surface electrodes (Ambu GmbH, Germany). Impedances were kept below 10 *k*Ω. After recording EEG and EMG, the signals were downsampled and saved at 2000 Hz.

The stimulation site targeting the right EDC was determined prior to each experimental session. For this, 40 stimuli were applied to the left motor cortex with an initial intensity of 40% maximum stimulator output (MSO). In case this intensity was insufficient to elicit MEPs, it was increased in steps of 5% MSO. At the three stimulation sites that resulted in the highest peak-to-peak amplitudes, three additional stimuli were applied. Of these, the stimulation site resulting in the most consistent and highest amplitude MEP was then considered as the motor hotspot. Next, the resting motor threshold (RMT) was determined, which was defined as the lowest intensity that resulted in at least 5 MEP out of 10 stimuli (23). Finally, we stimulated at seven different intensities (i.e., 90 – 150% RMT, in steps of 10% RMT) with 10 trials at each intensity. Blocks of the same intensity were applied in random order.

### Data cleaning and preprocessing

Data were imported into MATLAB R2022a (MathWorks, Natick, MA, USA) for preprocessing and analysis. Recordings from separate sessions were combined into single datasets for each participant, and stimulation artifacts were interpolated prior to bandpass filtering 1 - 40 Hz. For one participant, we were unable to include a subset of sessions due to data corruption issues. Following filtering, we used artifact subspace reduction (ASR) and independent component analysis (ICA) to remove artifacts. Finally, clean recordings were current source density transformed using the spherical spline method implemented with the publicly available SCD plugin for EEGLAB (24). For further details, see Supplement Methods.

### EMG and EEG Measures

Using EMG recorded from the right EDC, we computed MEP amplitudes as the log-transformed maximum peak-to-peak amplitudes within 15-50 ms after TMS stimulation. We also computed the standard deviation of the EMG signal from the right EDC for each trial. Spectral EEG measures were computed using Welch’s method and the FOOOF algorithm (25). which computed AE in the aperiodic model parameters. Next, to extract the alpha and beta phase of each EEG signal, we used the fast Fourier transform on Hanning windowed EEG signals. Finally, we applied two compression algorithms, CTW and LZ, to compute signal entropy. For further details, see Supplemental Methods.

### Statistical analysis

We used linear mixed models (LMMs) to predict log-scaled MEP amplitude with random intercepts for participants and fixed effects for the EEG feature of interest as well as pulse intensity (measured as percent MSO), pulse number, (i.e., the chronological index of the TMS pulse, which controls for habituation and facilitation effects), and the log-scaled standard deviation of the pre-pulse EMG recorded from the right EDC muscle (to control for muscle activation). All LMMs also accounted for interactions between pulse intensity and pulse number, as well as pulse intensity and muscle activation. To further assess the extent to which EEG phase features predicted CSE, we investigated the EEG channel with the smallest P-values for alpha phase and beta phase separately and then fit log-scaled and z- transformed MEP-amplitudes to a cosine function to compute modulation depth. To correct for multiple LMMs across 64 EEG channels, we utilized threshold free cluster enhancement (TFCE) (26) using parameters E = 2/3 and H = 2 as recommended by Mensen and Khatami (27) and nonparametric testing with 200 permutations. Finally, to assess the extent to which the predictive power of an EEG feature was driven by within-subject variance (i.e., a state marker) or between-subject variance (i.e., a trait marker), we derived ICCs and then took the quantity 1 - ICC to be high when a feature was a state marker and low when a feature was a trait marker (see Fig. 5). For further details, see Supplemental Methods.

## Results

ASR rejected a mean of 20.5 s of data across participants, and data from two participants were rejected due to an excessive number (> 10) of noisy channels. The 34 remaining participants had a mean of 6.7 noisy channels which were interpolated by ASR. See Fig. 1 for a summary of usable data in remaining participants. TMS trials were rejected based on MEP criteria, resulting in a mean of 45.1 trials rejected per participant due to the absence of a measurable or otherwise valid MEP (min = 5, max = 107, median = 42.5). The final clean dataset consisted of 8972 TMS trials across 34 participants, with a mean of 264 trials per participant (min = 45, max = 541, median = 237.5).

**Figure 1.**
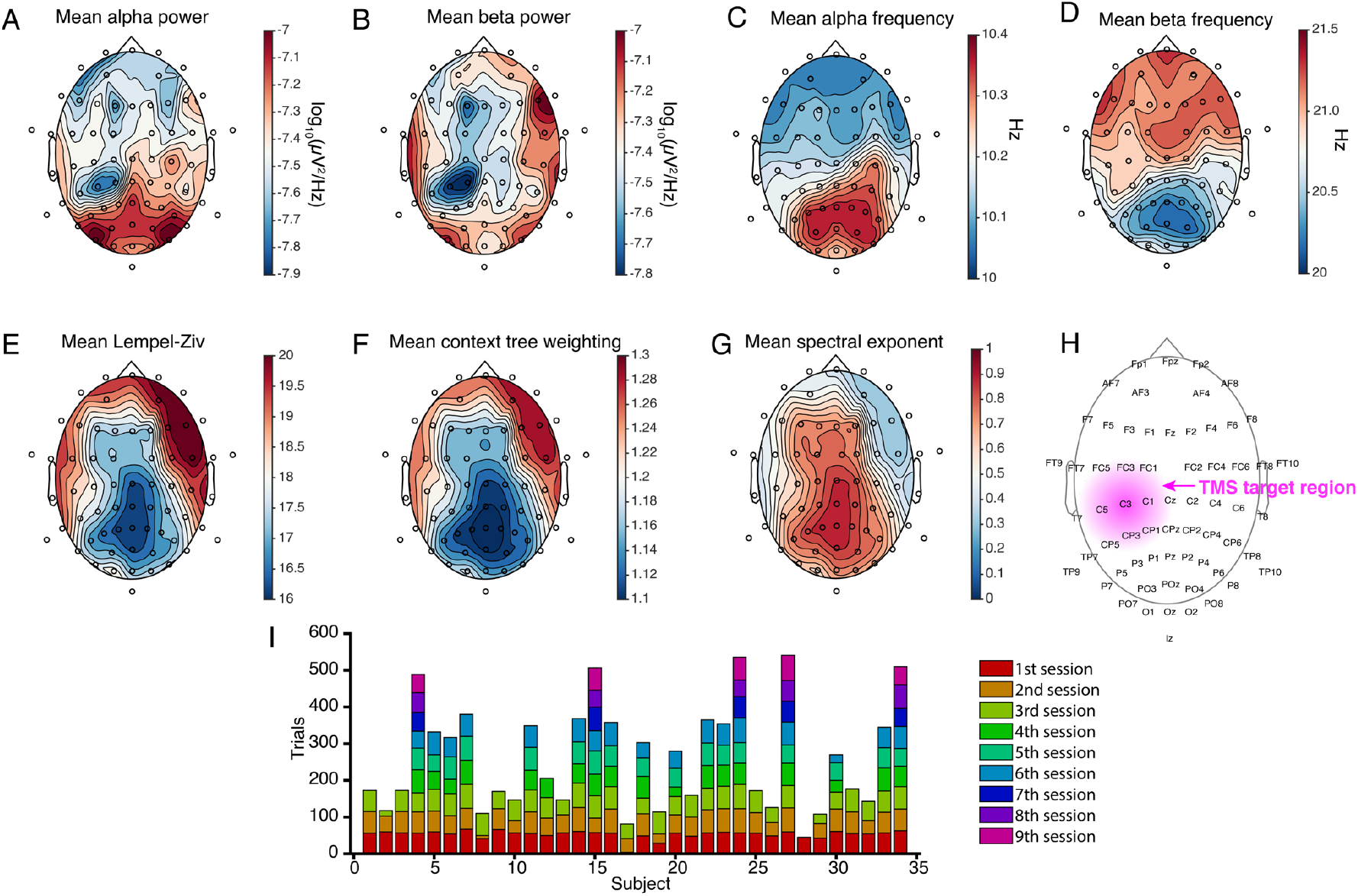
Topographic distributions of EEG features. Alpha power (A) shows a very different distribution than beta power (B). On the other hand, alpha frequency (C) and beta frequency (D) show similar topographies that appear to be mirror images of one another. BEMs (E-G) also show remarkably similar topographies with one another; entropy features (E,F) appear as the mirror image of the aperiodic exponent (G). EEG channel names are indicated in (H), as well as the TMS target region (magenta); note that this is an approximation and not based on real hotspot mapping data. Finally, the stacked bar plot in (I) provides an overview of how many TMS trials were taken from different sessions for each participant.

Having cleaned data, we estimated alpha and beta power, phase, and mean frequency, as well as BEMs CTW, LZ, and AE, for each participant, channel, and trial. We then visualized spatial distributions of relevant EEG features across the scalp (Fig. 1). As expected, EEG alpha power was highest over posterior regions and lowest over anterior regions (Fig. 1A). EEG beta power followed a different spatial distribution, with the highest power over peripheral regions (Fig. 1B). We observed a spatial minimum of both alpha and beta power at channel CP1, immediately posterior to the stimulation site, which is likely related to cumulative stimulation effects.

We also visualized the mean alpha and beta frequency across participants at each channel. By default, we defined the alpha band as 8 - 12 Hz and the beta band as 16 - 24 Hz, but using the fitting oscillations and one-over-f (FOOOF) algorithm (25), we adjusted these frequency ranges in a data-driven manner according to power spectral peaks for each channel of each participant’s EEG data (see Materials and Methods). If a spectral peak could not be identified, then the alpha and beta band ranges defaulted to those stated above. However, if a peak was identified near 10 Hz (alpha) or 20 Hz (beta), we took the peak frequency +/- 8 Hz as the frequency range, albeit with a limit imposed such that the range could not exceed 8 - 13 Hz (alpha) or 14 - 30 Hz (beta). This ensured that the frequency bands used in each case stayed within canonical limits. To visualize the results of this approach, we viewed topographies of the middle most frequency in the alpha (Fig. 1C) and beta (Fig. 1D) band, averaged across participants. The alpha band showed higher frequencies over posterior areas, whereas for the beta band, we observed the opposite sagittal gradient. Next, we visualized the spatial distribution of the broadband measures LZ, CTW, and AE, which all yielded remarkably similar topographies. LZ and CTW (Fig. 1E,F) were both minimal over parietal regions and maximal over frontotemporal regions, whereas AE (Fig. 1G) exhibited the same topography in reverse, with the steepest 1/f background evident over parietal regions and the least steep 1/f background evident over frontotemporal regions.

To visualize the overall strength of each EEG predictor, we first visualized t-statistcs from LMMs for each channel in the vicinity of the TMS target region. In the case of phase, we used the square root of the F-statistic as a proxy for the t-statistic, given that each phase model contained separate t-statistics for the sine and cosine of phase and, furthermore, that positive and negative values are not easily interpretable for circular variables such as phase. We plotted these values for central channels, demonstrating that beta band features (both power and phase) were generally more predictive than alpha band features (Fig. 2).

**Figure 2.**
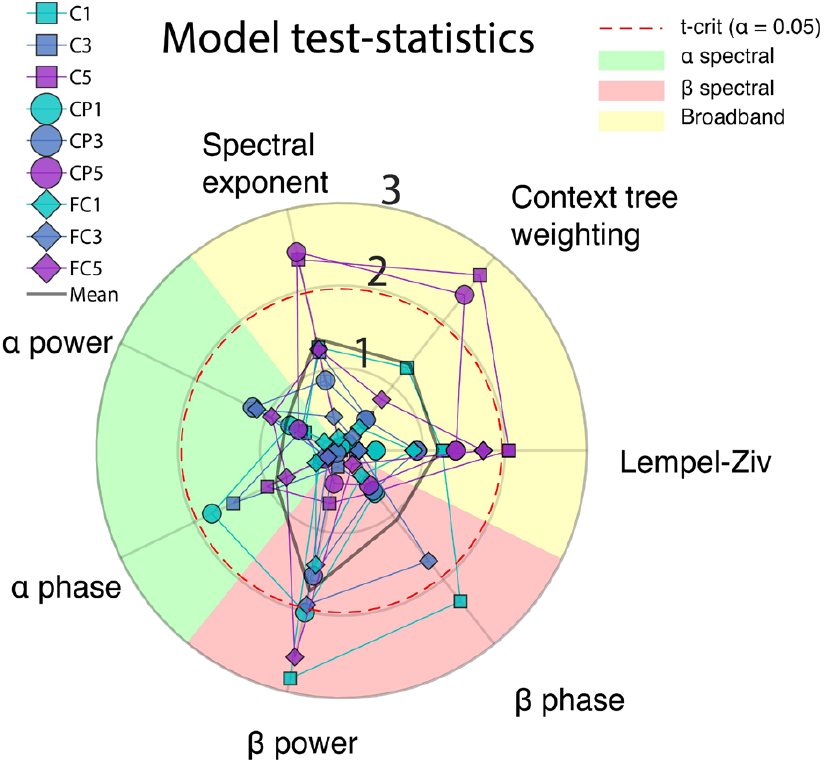
Exploratory visualization of EEG features in the proximity of the TMS target region. In an initial comparison of the predictive ability of different EEG features, we plotted the t-statistic (or *F*, in the case of phase) from 9 channels in the proximity of the TMS target region (left M1).

Using TFCE (26), which corrects for multiple comparisons while emphasizing channels that show similar effects as neighboring channels, combined with permutation testing to compute empirical P-values for each channel, we identified multiple channels for each bandlimited (Fig. 3) and broadband (Fig. 4) EEG feature that significantly predicted CSE. In what follows, all P-values from LMMs are reported as corrected using TFCE with permutation testing.

**Figure 3.**
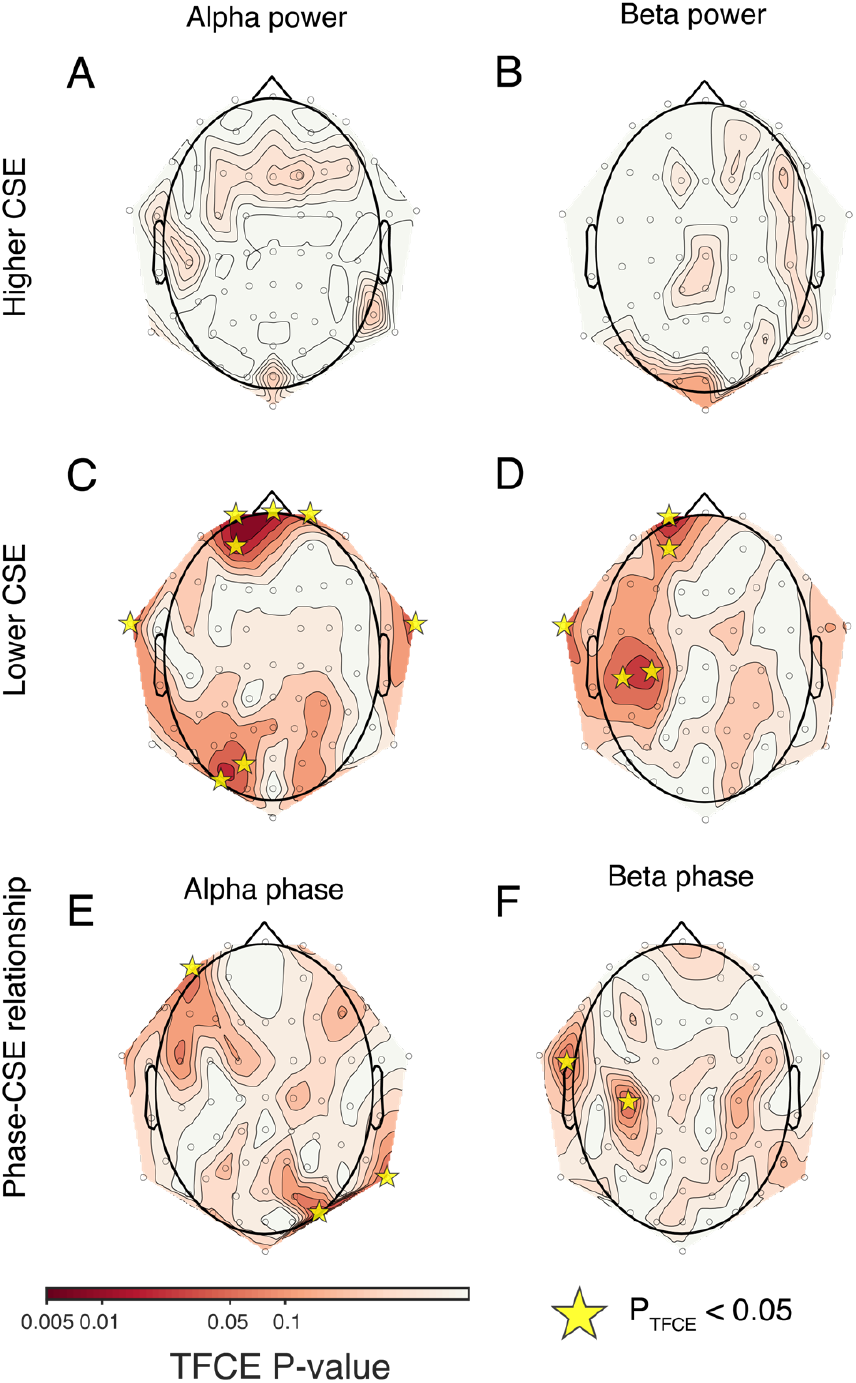
Topographic distributions of bandlimited spectral feature TFCE corrected P-values (linear mixed models). Higher EEG alpha and beta power were both associated with lower CSE; however, only beta power showed this effect in the vicinity of the TMS stimulation site. Note that, in the case of EEG alpha or beta phase, the test statistic used for clustering was the square root of the F-stat; thus, this test statistic always had a positive sign and thus the results which would be displayed in panel F and panel H are not applicable.

**Figure 4.**
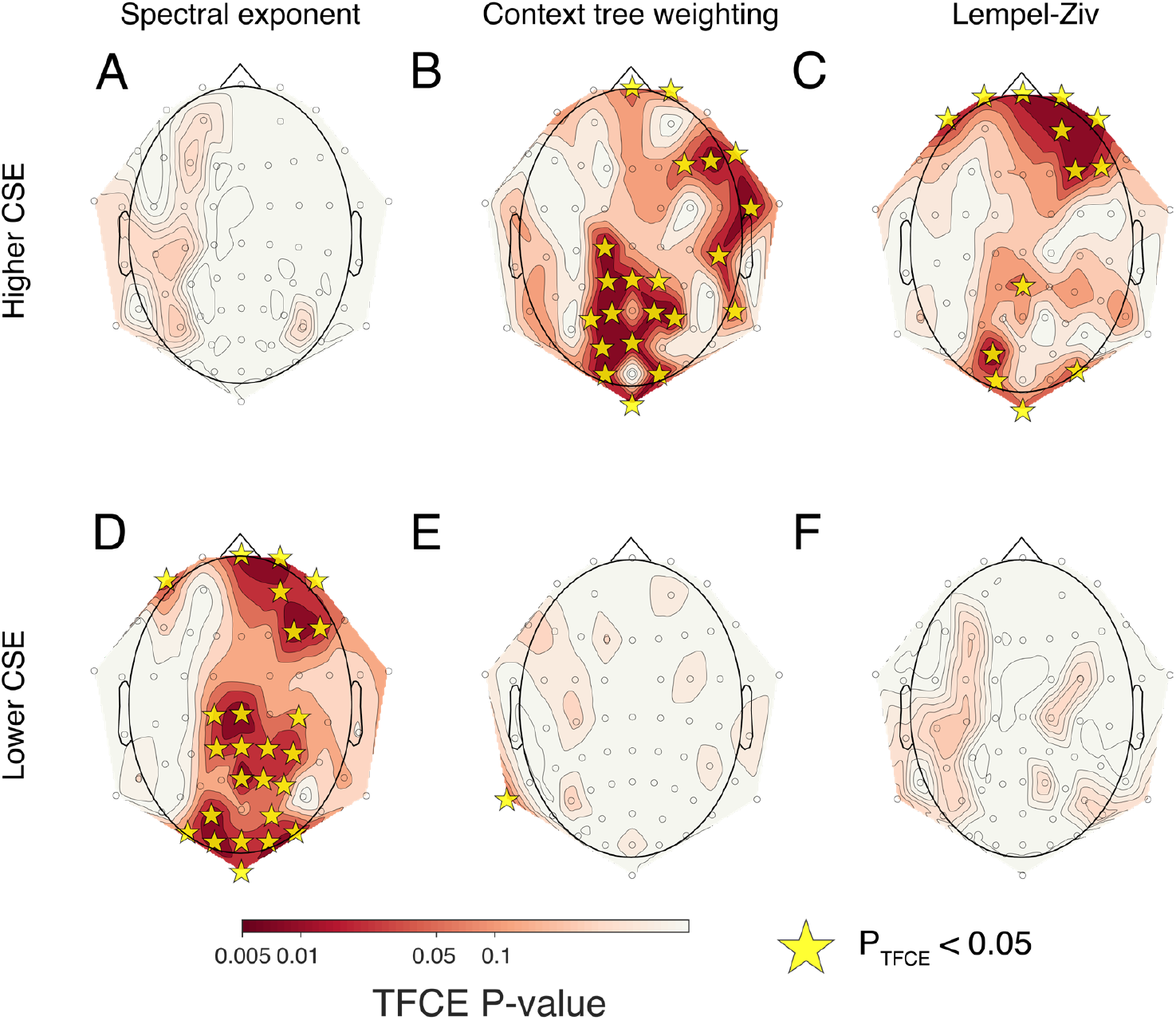
Topographic distributions of broadband feature TFCE corrected P-values (linear mixed models). EEG entropy was associated with higher CSE (Fig. 3B, C), whereas AE was associated with lower CSE (Fig. 3D). We obtained largely negative findings for increased CSE with AE (Fig. 1A) and decreased CSE with entropy (Fig. 1E,F).

First, we examined the predictive ability of bandlimited EEG features in the alpha and beta bands. In the case of both alpha and beta power, we found that decreases in power at some channels corresponded to significant increases in CSE. In the case of alpha power, this effect was largely prefrontal and thus distant from the TMS target site (Fig. 3C), whereas in the case of beta power, this effect was significant at a pair of prefrontal electrodes, but also a pair of electrodes centered at the TMS target region (C3, P_TFCE_ < 0.05, and C5, P_TFCE_ < 0.05, see Fig. 3D). In the case of EEG phase, alpha phase (Fig. 3E) was significantly predictive of CSE only at a number of largely peripheral channels. At the channel with the strongest predictive value as judged by LMMs, PO8, the modulation depth was not statistically different from randomly permuted data (P > 0.05). In the case of beta phase (Fig. 3F), two channels, C3 and FT7, showed a significant phase effect as judged by LMMs; of these channels, the more predictive of the two, C3 (P_TFCE_ < 0.05), was centered over the TMS target region. For exact TFCE- corrected P-values, see Table 1.

**Table 1.**
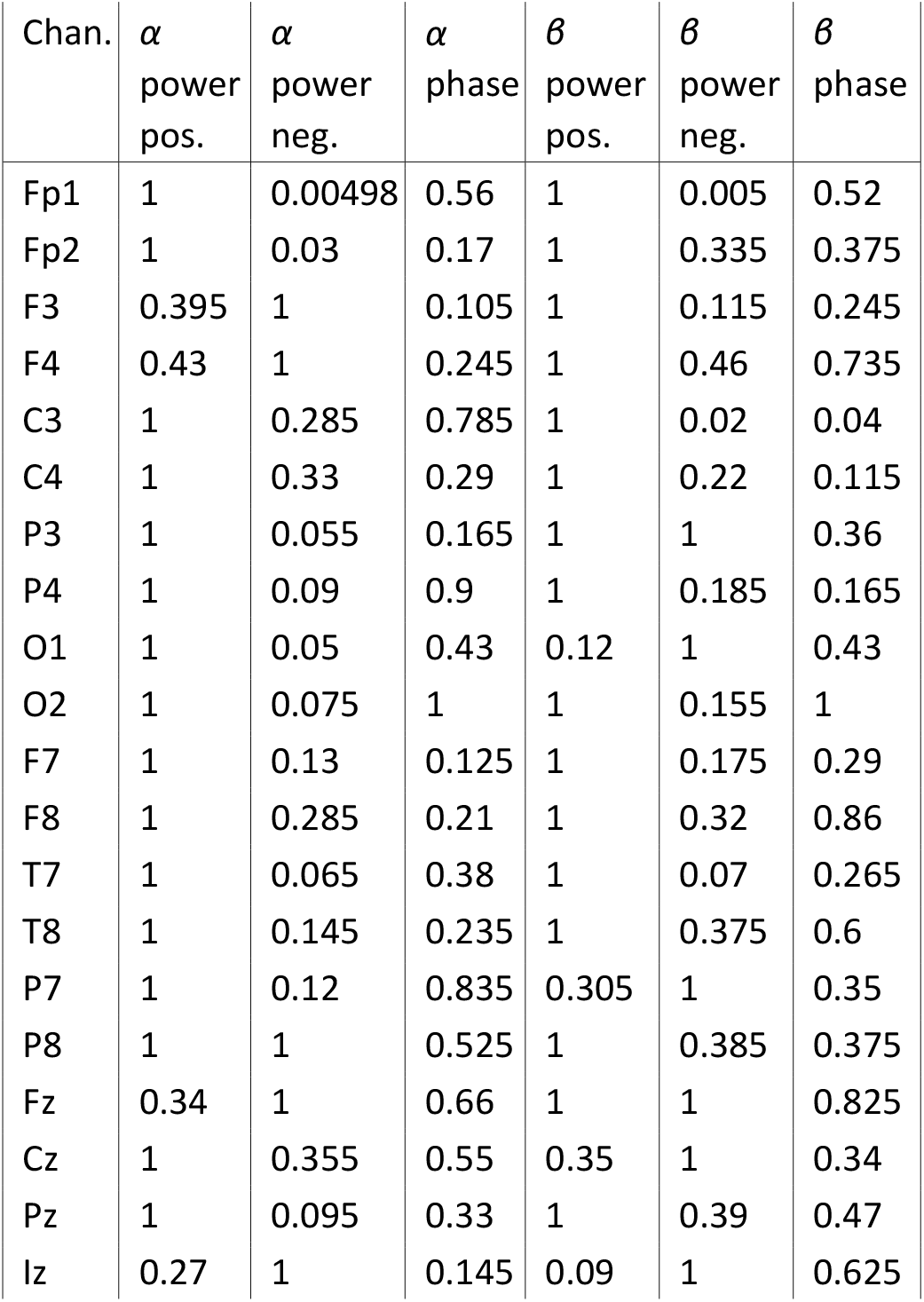
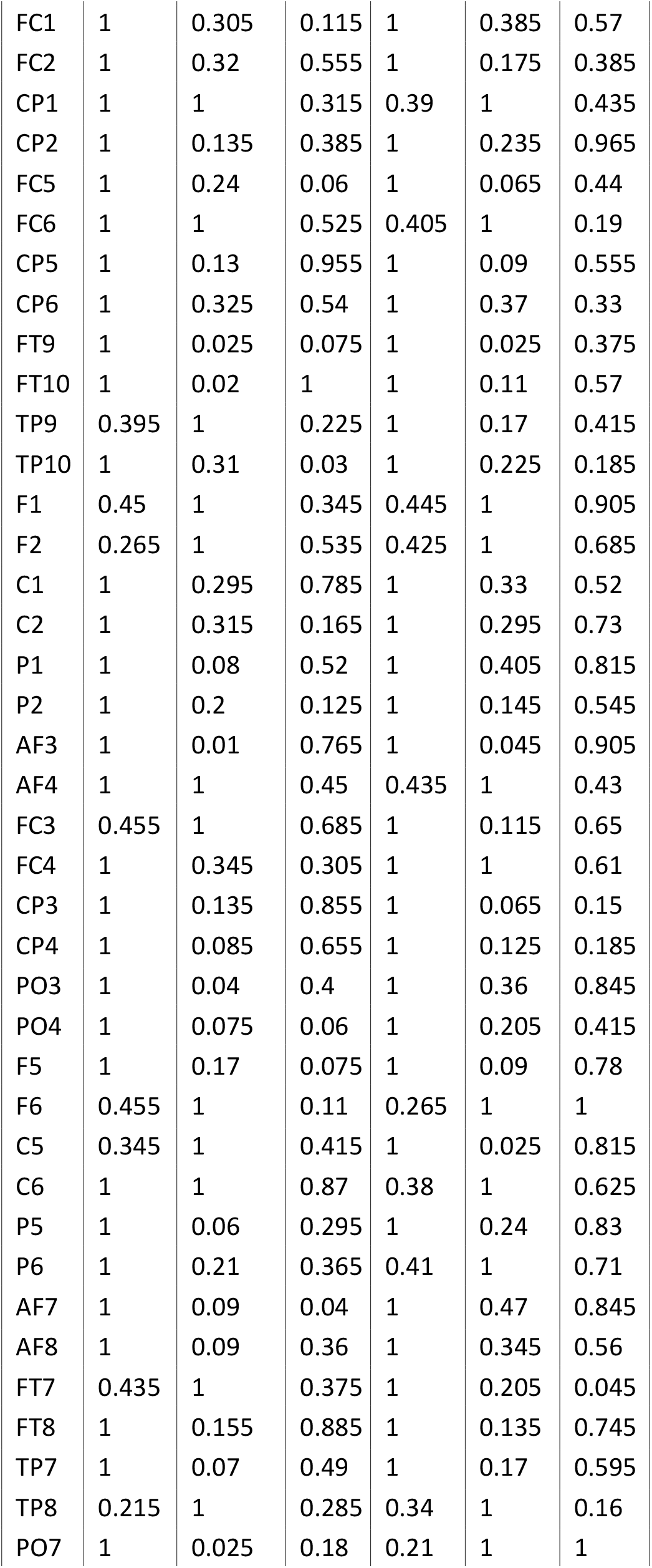
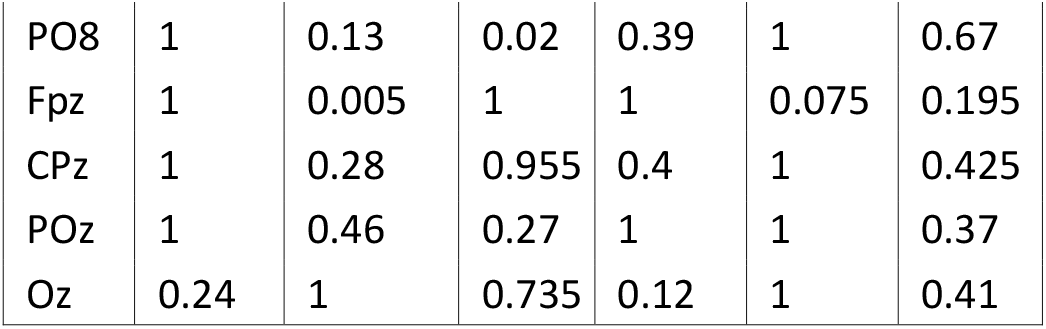
TFCE-corrected P-values from linear mixed models. Chan. = EEG channel, Pos. = positive effect, Neg. = negative effect.

Next, we examined the predictive ability of each BEM. Across a number of electrodes, we found that high AE corresponded to low CSE, whereas high CTW and high LZ corresponded to high CSE (Fig. 3); note one marginally significant peripheral channel (TP9) which showed the opposite effect for CTW and was likely a spurious finding (Fig. 3E). In general, all three measures showed significant effects both in prefrontal and centroparietal regions. However, only AE and CTW showed significant effects in the vicinity of left M1 targeted by TMS, specifically at channel C1 (Fig. 4B,D). Additionally, immediately adjacent posterior areas, including channels CP1 and CPz (Fig. 4B,D), also predicted CSE. These somatosensory regions are known to play a role in modulating motor cortex excitability (28–31). For exact TFCE-corrected P-values, see Table 2.

**Table 2.** TFCE-corrected P-values from linear mixed models. Chan. = EEG channel, Pos. = positive effect, Neg. = negative effect.

| Chan. | AE pos. | CTW pos. | LZ pos. | AE neg. | CTW neg. | LZ neg. |
| --- | --- | --- | --- | --- | --- | --- |
| Fp1 | 1 | 0.13 | 0.015 | 0.09 | 1 | 1 |
| Fp2 | 1 | 0.035 | 0.00498 | 0.005 | 1 | 1 |
| F3 | 0.32 | 0.395 | 1 | 1 | 1 | 0.27 |
| F4 | 1 | 0.025 | 0.005 | 0.005 | 1 | 1 |
| C3 | 0.255 | 1 | 1 | 1 | 0.29 | 0.25 |
| C4 | 1 | 0.135 | 0.19 | 0.045 | 1 | 1 |
| P3 | 1 | 0.015 | 0.08 | 0.09 | 1 | 1 |
| P4 | 1 | 0.005 | 0.28 | 0.03 | 1 | 1 |
| O1 | 1 | 0.00498 | 0.03 | 0.005 | 1 | 1 |
| O2 | 1 | 0.005 | 0.075 | 0.01 | 1 | 1 |
| F7 | 1 | 1 | 0.11 | 1 | 0.24 | 1 |
| F8 | 1 | 0.02 | 0.07 | 0.055 | 1 | 1 |
| T7 | 0.405 | 0.235 | 1 | 1 | 1 | 0.4 |
| T8 | 1 | 1 | 1 | 0.335 | 0.345 | 1 |
| P7 | 0.405 | 0.085 | 1 | 1 | 1 | 0.375 |
| P8 | 1 | 0.455 | 1 | 0.225 | 1 | 0.31 |
| Fz | 1 | 0.07 | 0.1 | 0.085 | 1 | 1 |
| Cz | 1 | 0.11 | 0.14 | 0.005 | 1 | 1 |
| Pz | 1 | 0.09 | 0.08 | 0.005 | 1 | 1 |
| Iz | 1 | 0.00498 | 0.045 | 0.03 | 1 | 1 |
| FC1 | 1 | 0.285 | 0.485 | 0.1 | 1 | 1 |
| FC2 | 1 | 0.07 | 0.1 | 0.105 | 1 | 1 |
| CP1 | 1 | 0.00498 | 0.055 | 0.015 | 1 | 1 |
| CP2 | 1 | 0.045 | 0.11 | 0.02 | 1 | 1 |
| FC5 | 1 | 1 | 0.33 | 1 | 0.49 | 1 |
| FC6 | 1 | 0.05 | 1 | 0.26 | 1 | 1 |
| CP5 | 0.305 | 0.235 | 1 | 1 | 1 | 0.285 |
| CP6 | 1 | 1 | 0.13 | 0.15 | 0.465 | 1 |
| FT9 | 0.44 | 1 | 0.16 | 1 | 0.39 | 1 |
| FT10 | 1 | 0.08 | 0.23 | 0.055 | 1 | 1 |
| TP9 | 0.365 | 1 | 1 | 1 | 0.02 | 0.195 |
| TP10 | 1 | 0.265 | 1 | 0.195 | 1 | 0.34 |
| F1 | 1 | 1 | 0.14 | 0.31 | 0.32 | 1 |
| F2 | 1 | 0.07 | 0.08 | 0.075 | 1 | 1 |
| C1 | 1 | 0.00498 | 0.1 | 0.02 | 1 | 1 |
| C2 | 1 | 0.255 | 1 | 0.055 | 1 | 0.305 |
| P1 | 1 | 0.005 | 0.265 | 0.11 | 1 | 1 |
| P2 | 1 | 0.00498 | 1 | 0.02 | 1 | 0.375 |
| AF3 | 0.43 | 0.4 | 0.175 | 1 | 1 | 1 |
| AF4 | 1 | 1 | 0.005 | 0.01 | 0.49 | 1 |
| FC3 | 0.5 | 1 | 1 | 1 | 0.385 | 0.28 |
| FC4 | 1 | 1 | 1 | 0.14 | 0.395 | 0.34 |
| CP3 | 1 | 0.375 | 0.375 | 0.335 | 1 | 1 |
| CP4 | 1 | 0.13 | 0.055 | 0.01 | 1 | 1 |
| PO3 | 1 | 0.00498 | 0.00498 | 0.005 | 1 | 1 |
| PO4 | 1 | 0.225 | 0.41 | 0.03 | 1 | 1 |
| F5 | 1 | 1 | 0.225 | 0.245 | 0.35 | 1 |
| F6 | 1 | 0.005 | 0.02 | 0.01 | 1 | 1 |
| C5 | 0.27 | 0.14 | 1 | 1 | 1 | 0.235 |
| C6 | 1 | 0.01 | 0.32 | 0.125 | 1 | 1 |
| P5 | 0.22 | 1 | 1 | 1 | 0.23 | 0.335 |
| P6 | 0.335 | 0.47 | 1 | 1 | 1 | 0.36 |
| AF7 | 1 | 0.05 | 0.005 | 0.01 | 1 | 1 |
| AF8 | 1 | 0.15 | 0.005 | 0.035 | 1 | 1 |
| FT7 | 0.385 | 0.115 | 1 | 1 | 1 | 1 |
| FT8 | 1 | 0.005 | 0.31 | 0.195 | 1 | 1 |
| TP7 | 1 | 0.07 | 0.24 | 0.32 | 1 | 1 |
| TP8 | 1 | 0.025 | 0.065 | 0.115 | 1 | 1 |
| PO7 | 1 | 0.065 | 0.06 | 0.035 | 1 | 1 |
| PO8 | 1 | 0.15 | 0.03 | 0.01 | 1 | 1 |
| Fpz | 1 | 0.03 | 0.00498 | 0.00498 | 1 | 1 |
| CPz | 1 | 0.01 | 0.04 | 0.015 | 1 | 1 |
| POz | 1 | 0.00498 | 0.345 | 0.06 | 1 | 1 |
| Oz | 1 | 1 | 0.395 | 0.015 | 0.42 | 1 |

Finally, to determine whether band-limited features and BEMs were primarily trait markers explaining between-subjects variance or state markers explaining within-subject variance, we computed the intraclass correlation (ICC) for AE, CTW, LZ, alpha power, and beta power. We did not compute ICCs for alpha or beta phase, as oscillatory phase fluctuates many times per second and is thus a state marker by definition. The ICC is high when values from within a subject show strong agreement indicating low within-subject variance and suggesting a relatively stable or trait marker. Conversely, the ICC is low when within-subject values show weak agreement, reflecting high within-subject variance and suggesting a more dynamic or state marker. To capture this trait-state continuum, we calculated the quantity 1 - ICC (Fig. 5).

BEMs AE, CTW, and LZ generally fell closer to the state end of the continuum at most locations (Fig. 5A-C), whereas spectral alpha and beta power generally fell closer to the trait end of the continuum (Fig. 5D,E). The trait-like tendency of spectral power features and state-like tendency of broadband features were also revealed by histograms (Fig.5F). Overall, CTW appeared to be the most state-like measure. Taken together, the above findings demonstrate that BEMs are relatively dynamic and thus likely to be more sensitive to cortical state changes.

**Figure 5.**
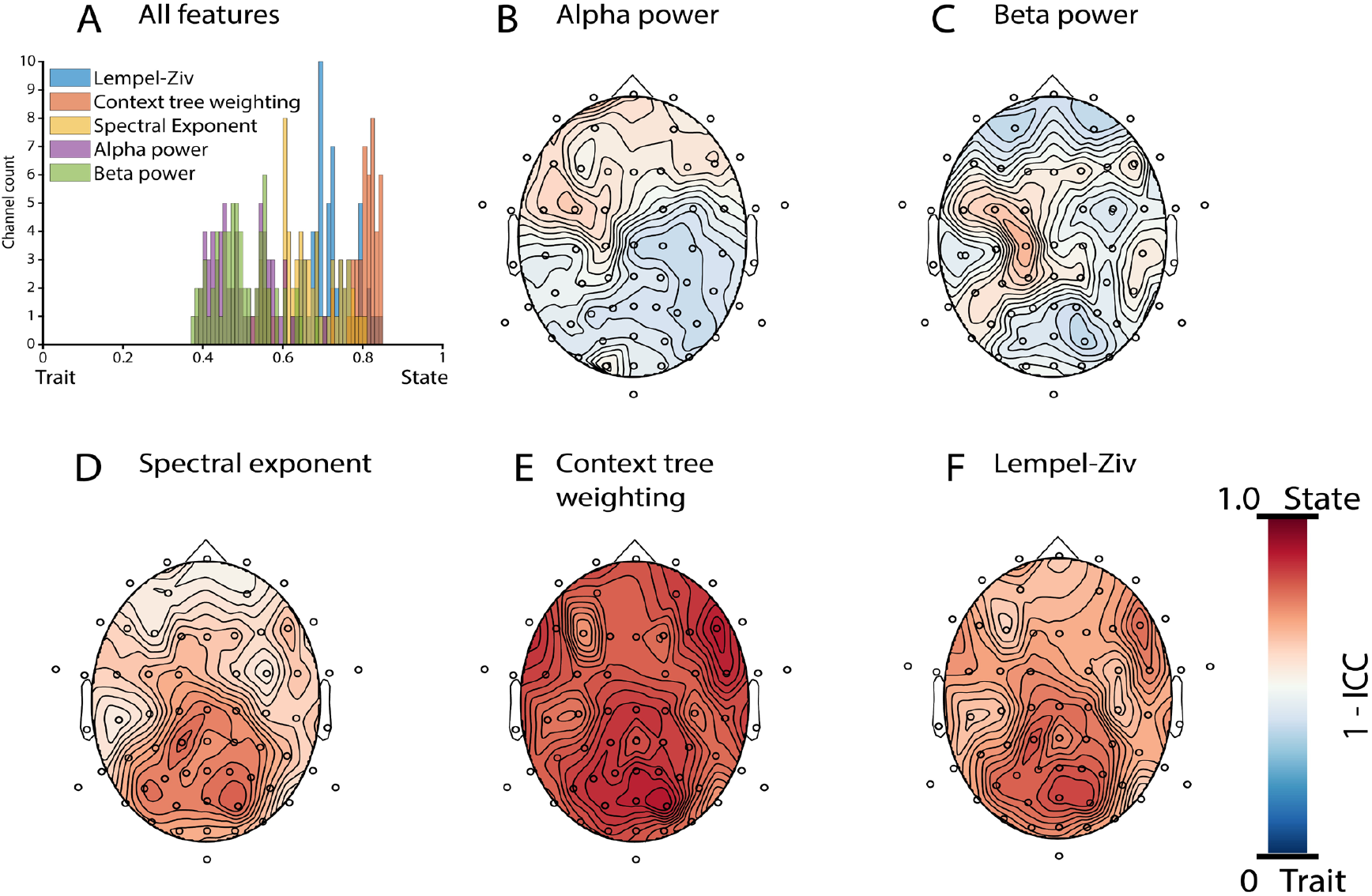
State versus trait features. As judged by the ICC, BEMs appear to be state measures (i.e., more variance within subjects than between subjects, Fig. 5D-F), whereas spectral power EEG features appear to be trait measures (i.e., more variance between subjects than within subjects, Fig. 5B,C). To measure the “stateness” of a feature, we subtracted the intraclass correlation (ICC) from 1, yielding a quantity that is greatest for state measures (red colors in topoplots) and lowest for trait measures (blue colors in topoplots). A histogram of this quantity (Fig. 5A) shows the number of channels for each measure which fall into bins along the trait-state continuum.

## Discussion

In this study, we utilized the aperiodic exponent (AE) and entropy measures (CTW, LZ) derived from broadband EEG signals to predict CSE during TMS pulses applied independently of brain state in healthy participants. Our findings indicate that BEMs (AE, CTW, and LZ) are significant predictors of CSE (Fig. 4), comparable to conventional band-limited EEG features (Fig. 3). Among the band-limited features, beta power (Fig. 3D) and phase (Fig. 3F) demonstrated the highest topographical specificity as a marker for motor CSE. By contrast, AE, CTW, and LZ were more sensitive to variations in topographically diffuse brain states (Fig. 4), which appeared to vary more within subjects than alpha or beta power (Fig. 5). These broadband measures, characterized by lower ICCs, were more indicative of dynamic state changes, making them particularly suitable for identifying moments of widespread brain excitability. Thus, AE, CTW, and LZ could be useful for marking states of brain excitability beyond just the motor cortex. Furthermore, the BEMs we evaluated were measured from roughly 1 s time windows and varied on a slower timescale than EEG phase features which, by definition, fluctuate many times per second. As such, AE, CTW, and LZ might be less susceptible to brief physiological artifacts which influence phase estimates.

### BEMs predict CSE

In response to the challenges of bandlimited features for CSE prediction, AE, CTW, and LZ eliminate issues stemming from the experimenter’s choice of EEG frequency and/or phase. These BEMs may encompass multiple EEG frequency bands, thus avoiding the pitfalls of a bandlimited approach. Two of these BEMs, CTW and LZ, are based on eponymous compression algorithms which quantify the number of unique substrings or states in the signal (32, 33). The other BEM, AE, is based on the spectral steepness of the 1/f background of the EEG signal, which has been used a proxy for the ratio of neural excitation to inhibition (21, 34). We found that BEMs significantly predicted CSE, albeit in slightly different regions in each case. AE and CTW significantly improved the model fit in the vicinity of the left M1 which we targeted with TMS (Fig. B,D), including channel C1, as well as left centroparietal channels, fitting with another prior study of surface-Laplacian transformed EEG signals (35) which reported that postcentral recordings can be used to predict CSE in a posthoc analysis of data collected during TMS of left M1.

### Different EEG entropy features offer different advantages

Remarkably, LZ and CTW show quite different sensitivities as entropy estimators, despite both being computed from compression algorithms. Why is LZ, unlike CTW, mainly useful as a CSE predictor when computed from prefrontal electrodes? While it is plausible that CSE is influenced by prefrontal disinhibition of motor cortex (e.g., executive control), frontal electrodes may have been contaminated by residual muscle artifacts to a greater extent than centroparietal channels, despite our best efforts to reduce physiological artifacts from EEG signals and to covary for prepulse muscle tone recorded from the right hand. Thus, we are less confident that the LZ finding is truly a neural effect, as opposed to a readout of facial muscle activation. CTW is perhaps a more useful measure of cortical entropy than LZ for the purpose of inferring CSE. This is somewhat surprising, given that both entropy estimates are based on compression algorithms and have performed similarly on neural data in other studies (19, 36–38), and, furthermore, LZ is much more widely applied in human neuroscience across contexts ranging from studies of anesthetics (11, 12, 15, 39), sleep (12, 14, 16, 40), and pharmacological drug challenge (14, 41, 42), including effects on TMS-evoked potentials (18, 43, 44). Nonetheless, from a purely computational perspective, there are strong reasons to believe CTW is a superior entropy estimator for time series data. Unlike LZ, CTW is based on Bayesian inference, which has long been known to produce more data-efficient estimators (45). Accordingly, CTW consistently outperforms other entropy estimators (including LZ) when evaluated on synthetic data (46). This increased data efficiency allowed us to discretize the EEG signal into 8 symbols prior to the calculation of CTW (compared to the LZ calculation with only 2 symbols), such that CTW was able to capture more details of the signal’s fine-grained dynamics. Considering the above, we advocate for further use of CTW as an entropy estimator for neural data, not only in TMS-EEG experiments, but also in other EEG analysis contexts.

### A new approach for state-dependent TMS

We carried out this study with an eye toward state-dependent TMS applications, e.g., closed-loop TMS, guided by EEG features. As suggested by its name, TMS in this context is automatically triggered by variables which indicate a rapidly fluctuating cortical state. Variables which primarily reflect a trait, rather than dynamic state, are not expected to be useful for state-dependent TMS, given that they are approximately static at the relevant timescale within a given subject.

Given the hierarchical nature of our data, which included many TMS-EEG trials both within and between participants, it was reasonable for us to ask whether our findings were driven primarily by state (within-subject variance) or trait (between-subjects variance). Although the entropy (12–15, 19) and 1/f background (21, 47, 48) of the broadband EEG is already known to change drastically between states of consciousness, it is currently uncertain to what extent these EEG features change within the awake and alert resting state. To address this uncertainty, we computed ICCs for EEG features and used the quantity 1−ICC to infer the degree of within-subject variance, with 1 indicating a state variable and 0 indicating a trait variable. We found that BEMs generally showed greater within-subject variance than spectral alpha or beta power. Our results suggest that BEMs are sensitive to cortical states that fluctuate within a TMS-EEG session.

### Conclusions, limitations, and future directions

Hitherto, the field of state-dependent TMS has focused almost exclusively on bandlimited spectral EEG measures such as alpha and beta power or phase. Our results suggest the utility of new, and potentially complimentary, BEMs that also predict CSE. Because these new EEG measures do not require narrowband filtering nor phase estimation/prediction, they are unaffected by many issues which have likely hampered the reproducibility of previous work in this field. At the same time, we admit that the BEMs we tested are not parameter free, e.g., other researchers may choose different FOOOF parameters (25) when computing AE, thus introducing further researcher degrees of freedom (49). However, we maintain that these choices are more tractable than the current methodological divergences which impede the field. Furthermore, although our first pass at data cleaning was handled using an automated procedure with objective criteria (ASR), we also utilized a group-level ICA decomposition for further cleaning (e.g., to remove technical artifacts). Because the ICA decomposition is generally slow and thus cannot be run in real time, this step needs to be replaced in the future to make our results translatable to real-time applications. We tested BEMs only in young, healthy participants, and future work will need to extend this analysis to clinical cohorts to assess their utility in therapeutic applications. Once validated in clinical populations, the practical value of these EEG features as dynamic CSE predictors will be tested in state-dependent TMS-EEG. Ultimately, this line of research aims to use EEG entropy and the 1/f background as TMS triggers in closed-loop experiments.

In conclusion, this study identifies band-limited and broadband features as complementary markers of CSE. Beta-band activity emerges as a spatially localized marker, topographically specific to the motor cortex. The oscillatory beta phase, which fluctuates several times per second, inherently serves as a state marker of CSE, while low beta power corresponds to high CSE and is primarily a trait-like marker explaining between-subject variance. In contrast, entropy and aperiodicity provide valuable complementary insights as global brain state markers. These broadband features, which fluctuate on a slower timescale than oscillatory phase – seconds rather than milliseconds – provide potentially more robust brain state markers explaining within-subject variance, with entropy associated with high CSE and the aperiodic exponent associated with low CSE. These findings hold promise for advancing state-dependent TMS applications, particularly in therapeutic contexts.

## Acknowledgements

We sincerely thank all participants who volunteered for our study. We gratefully acknowledge the German Federal Ministry of Education and Research (BMBF) grant Bevares (13GW0570), and the European Union’s Joint Programme for Neurodegenerative Disease Research (EU-JPND 2022-130) grant Recast (01ED2309). We also gratefully acknowledge support from the Open Access Publishing Fund of the University of Tübingen.

## Author contributions

JF, SR, and AG conceived the idea for the project. JF wrote the analysis code, analyzed the data, and wrote the first draft of the manuscript. SR guided the analyses and advised the statistical analysis. BT assisted with data collection, as well as MEP data preprocessing and trial rejection. MK advised the analysis and performed the code review. PM advised with the use of entropy measures and provided the code to compute entropy features. AG supervised the project. All authors revised and approved the manuscript. Additionally, the manuscript was refined for grammatical accuracy, sentence structure and wording using a convolutional neural network-based tool (DeepL) and a large language model (ChatGPT).

