## Supplemental Methods for "Brain signal complexity and aperiodicity predict human corticospinal excitability"

**Data cleaning and preprocessing.** Stimulation trigger annotations were corrected to adjust for a 6 ms delay. Stimulation artifacts in EEG and EMG recordings were then interpolated prior to filtering. This was accomplished by 1) replacing the stimulation artifact (50.5 ms duration) with Not-a-Number (NaN) elements, 2) applying a 75 ms moving average filter to smooth a copy of the signal (MATLAB: movmean with 'omitnan' option), and 3) replacing NaN elements of the original, unsmoothed signal with the corresponding section from the smoothed signal. Next, filters were implemented using finite-impulse response (FIR) filtering (filter order = 8000, i.e., 4 x sampling rate). EEG recordings were then bandpass filtered 1 - 40 Hz (EEG), and EMG recordings were first notch filtered at 50 Hz (to attenuate linenoise) and then bandpass filtered 4 - 100 Hz.

Following filtering, EEG data were cleaned using ASR with the following parameters: ChannelCriterion = 0.7, LineNoiseCriterion = 4, BurstCriterion = 3, WindowCriterion = 0.3, ChannelCriterionMaxBadTime = 0.5, BurstRejection = 'off', Highpass = 'off'. ASR was chosen for this cleaning step because it can be used in real-time situations (1), giving findings from this experiment greater translatability to state-dependent TMS applications. Channels marked bad by ASR were spherically interpolated. Next, data were epoched 1000 ms pre-pulse to 200 ms post-pulse and baseline corrected. Finally, to further clean data, a group-level ICA decomposition was performed using the fastica algorithm on all concatenated EEG data downsampled to 125 Hz. Following the group-level ICA decomposition, we removed one component corresponding to a blink artifact, one component corresponding to a saccade artifact, four components corresponding to muscle artifacts, and eight components possibly corresponding to mechanical artifacts resulting from contact between the TMS coil and electrodes.

**Spectral EEG measures.** All spectral measures were computed using the FOOOF algorithm by (2) . For each EEG channel and trial computed the power spectral density (PSD) using the EEG signal from 1000 to 50 ms prior to stimulus onset with the pwelch() function in MATLAB [input parameters: WINDOW = 512, NOVERLAP = 256, F = linspace(1,40,100)]. We then modeled the PSD between 1 and 40 Hz using FOOOF with the 'fixed' approach to parameterizing the aperiodic component (i.e., the 1/f background) and 0.6 and 1.0 Hz as the lower and upper bounds, respectively, for fitted peak widths.

Having modeled the EEG PSD using FOOOF, AE was returned in the aperiodic model parameters. This measure, sometimes also referred to as the power law exponent, spectral slope, or spectral exponent, is the slope of the aperiodic component modeled by FOOOF, which is the EEG 1/f background, so named because its power is inversely related to frequency (3). When frequency and power are plotted in logarithmic coordinates and spectral peaks are subtracted, the 1/f background shows a linear relationship with frequency (4). It is then straightforward to compute AE as the slope of the 1/f background. Because this slope is nearly always negative, AE is generally reported as an absolute value. Evidence from a variety of experiments, including propofol in monkeys and theta modulated excitability in rat hippocampus (5), demonstrates that AE (steepness) increases with greater neural inhibition and decreases with greater neural excitation. As such, we hypothesized that low AE measured from motor cortical EEG signals should predict high CSE.

To identify the alpha peak frequency in a given trial and channel, we then looked for oscillatory components returned by FOOOF with peak frequencies between 8 and 13 Hz. If more than one oscillatory component with a peak frequency in this range was returned, we used the component with the peak frequency closest to 10 Hz. The alpha band was then defined for that trial and channel as $f$ +/- 2 Hz, where $f$ is the peak frequency. In the case that no oscillatory component was returned in the alpha range, we defined the alpha band as 8 - 12 Hz.

Similarly, to identify the beta peak frequency in a given trial and channel, we looked for oscillatory components returned by FOOOF with peak frequencies between 14 and 30 Hz. If more than one oscillatory component was returned in this range, we used the component with the peak frequency closest to 20 Hz. The beta band was then defined for that trial and channel as $f$ +/- 4 Hz, where $f$ is the peak frequency. If no oscillatory component was returned in the beta range, we defined the beta band as 16 - 24 Hz.

Having defined the alpha and beta band for the trial and channel under consideration, to extract alpha and beta power, we then computed the PSD of each filtered signal using the pwelch() function [input parameteres: F = linspace(0,30,128) and default values for WINDOW and NOVERLAP]. Alpha and beta power were computed using trapezoidal integration of the PSD between the appropriate limits of integration for each band, and this power estimate was then log-scaled. Finally, to extract the alpha and beta phase of each EEG signal, we applied the fast Fourier transform to a Hanning windowed segment of signal corresponding to twice the period of the trial-specific alpha or beta center frequency (6) taken from the end of the signal, stopping 50 ms before the TMS pulse. For example, if the EEG alpha band was determined by FOOOF as 9 - 13 Hz, we applied FFT to the signal from -232 to -50 ms relative to the TMS pulse, based on a center alpha frequency of 11 Hz and period of 90.9 ms. We then extracted the phase of the Fourier spectrum corresponding to the alpha or beta center frequency (e.g., in the example given above, the 11 Hz alpha phase). Next, we adjusted the phase estimate to correct for the fact that the FFT window ended 50 ms before the TMS pulse (e.g., if the frequency extracted by FOOOF was 10 Hz, then we added 180° to the phase estimate).

**Entropy analysis .** Entropy is commonly defined as the number of possible configurations of a system (e.g., thermodynamic entropy). A common approach to estimating signal entropy is to apply a compression algorithm to the signal following a symbolic transformation (e.g., discretization using quantiles). More complex or diverse signals are harder to compress, as they contain more information. Indeed, entropy and information are intimately related and, in some frameworks (7, 8), identical. We estimated signal entropy for each EEG channel and trial using the EEG signal from 1000 to 50 ms prior to stimulus onset (i.e., 0.95 s of data). To speed up computations, signals were first downsampled to 200 Hz. CTW entropy was computed by transforming each EEG signal into discrete octiles and then applying the CTW compression algorithm (9) to estimate the entropy rate (10). LZ entropy was computed by transforming the signal into two discrete states based on a median split of time series values. We then applied the LZ76 algorithm (i.e., the first in a series of Lempel-Ziv compression algorithm published in 1976) to estimate the signal's entropy (11). Because all trials had the same signal length, we did not normalize the entropy estimate according to signal length.

We used linear mixed models (LMMs) to predict log-scaled MEP amplitude with random intercepts for participants and fixed effects for the EEG feature of interest as well as pulse intensity (measured as percent MSO), pulse number, (i.e., the chronological index of the TMS pulse, which controls for habituation and facilitation effects), and the log-scaled standard deviation of the pre-pulse EMG recorded from the right EDC muscle (to control for muscle activation). All LMMs also accounted for interactions between pulse intensity and pulse number, as well as pulse intensity and muscle activation.

To test $K$ predictors of CSE in data containing $N$ trials across $n$ participants, we used separate LMMs at each channel with the formula

$$y = X\beta + Zu + \epsilon \quad [1]$$

where $y$ is the $N$ x 1 vector of log-scaled MEP amplitudes (i.e., proxies for CSE) for each trial, $X$ is the $N$ x $K$ matrix of fixed effects, including a fixed intercept term (first column), the EEG predictor (second column), the pulse intensity (third column), the pulse number (fourth column), the log-scaled standard deviation of the prepulse EMG signal (fifth column), and interactions of fixed effect terms (subsequent columns); $\beta$ is the $K$ x 1 vector of fixed effect coefficients, $Z$ is the $N$ x $n$ design matrix of random effects, $u$ is the $n$ x 1 vector of random intercepts, and $\epsilon$ is the $N$ x 1 vector of residuals. Note that in cases where the alpha or beta phase was used to predict CSE, we included two separate columns for the EEG predictor, one corresponding to the sine of the phase and the other corresponding to the cosine of the phase.

To further assess the extent to which EEG phase features predicted CSE, we investigated the EEG channel with the smallest P-values for alpha phase and beta phase seperately and then fit log-scaled and z-transformed MEP-amplitudes to a cosine function [as done previously, e.g. see (12)] with the formula

$$M = A \cdot \cos(\theta + \phi) + \epsilon \quad [2]$$

where $M$ is the vector of log-transformed and z-scored MEP amplitudes, $A$ is the modulation depth, $\theta$ is the oscillatory phase preceding the TMS pulse, $\phi$ is the optimal phase offset, and $\epsilon$ is the error term. Because the $M$ was already z-scored and thus mean-centered, there was no need to include a vertical offset in the model. We then tested the cosine fit using a permutation test in which the data were randomly shuffled 1000 times. An empirical P-value was computed by comparing the actual modulation depth to the surrogate distribution.

To correct for multiple LMMs across 64 EEG channels, we utilized threshold free cluster enhancement (TFCE) (13) using parameters E = 2/3 and H = 2 as recommended by Mensen and Khatami (14). This approach privileges channels surrounded by neighboring channels showing similar effects yet departs from previous cluster statistical approaches (15) by taking many different thresholds into consideration, rather than a single arbitrary threshold which determines clusters. In our case, we sampled 50 evenly spaced thresholds for test statistics starting from 0 and increasing to the largest test statistic present in the data (as judged by absolute value). We then used permutation testing to derive an empirical P-value for each electrode. Data were permuted by randomly shuffling the outcome variable, i.e., MEP amplitudes, for each of 200 permutations. For all thresholds explored, channels were considered neighbors if they fell within 55 mm of one another in the Euclidian space of a 3-dimensional standard template.

To assess the extent to which the predictive power of an EEG feature was driven by within-subject variance (i.e., a state marker) or between-subject variance (i.e., a trait marker), we derived ICCs according to the definition "ICC(l) for Case 1" given in McGraw and Wong (16) with publicly available code (17). We then took the quantity 1 - ICC to be high when a feature was a state marker and low when a feature was a trait marker (see Fig. 5).
